# Engraftment, fate, and function of HoxB8-conditional neutrophil progenitors in the unconditioned murine host

**DOI:** 10.1101/2021.12.20.473543

**Authors:** Joshua T. Cohen, Michael Danise, Kristina D. Hinman, Brittany M. Neumann, Renita Johnson, Zachary S. Wilson, Anna Chorzalska, Patrycja M. Dubielecka, Craig T. Lefort

**Author notes:** Corresponding Author: Craig T. Lefort, PhD, Rhode Island Hospital, 593 Eddy Street, Providence, RI 02903. These authors contributed equally (Joshua T. Cohen, Michael Danise).

## Abstract

The development and use of murine myeloid progenitor cell lines that are conditionally immortalized through expression of HoxB8 has provided a valuable tool for studies of neutrophil biology. Recent work has extended the utility of HoxB8-conditional progenitors to the *in vivo* setting via their transplantation into irradiated mice. Here, we describe the isolation of HoxB8-conditional progenitor cell lines that are unique in their ability to engraft in the naïve host in the absence of conditioning of the hematopoietic niche. Our results indicate that HoxB8-conditional progenitors engraft in a β1 integrin-dependent manner and transiently generate donor-derived mature neutrophils. Furthermore, we show that neutrophils derived *in vivo* from transplanted HoxB8-conditional progenitors are mobilized to the periphery and recruited to sites of inflammation in a manner that depends on the C-X-C chemokine receptor 2 and β2 integrins, the same mechanisms that have been described for recruitment of endogenous primary neutrophils. Together, our studies advance the understanding of HoxB8-conditional neutrophil progenitors and describe an innovative tool that, by virtue of its ability to engraft in the naïve host, will facilitate mechanistic *in vivo* experimentation on neutrophils.

## Introduction

As mediators of the acute inflammatory response, neutrophils play essential roles in host defense against bacterial and fungal infection (Ley et al., 2018). Neutrophil development into their mature effector state primarily occurs in the bone marrow. Neutrophils are abundant and short-lived, a combination that requires the expenditure of a great amount of energy and resources to produce around a billion mature neutrophils per kilogram human body mass per day under homeostasis (Kubes, 2018). The rationale for the host’s investment in the neutrophil is underscored by the manifestation of recurrent and severe infections in individuals with neutropenia or neutrophil dysfunction (Ley et al., 2018).

While primary neutrophils are easily obtained from blood or tissue samples, they are non-proliferative and have a short lifespan. Moreover, common techniques for introducing genetic material to cells result in artefactual activation of primary neutrophils. Thus, even in the current age of facile genetic engineering, mechanistic investigation of neutrophil biology remains a challenge. For decades, neutrophil biologists have relied on imperfect “neutrophil-like” cell lines that fail to fully recapitulate the breadth of neutrophil functionality and may differ from primary neutrophils mechanistically in those functions. Such models include human HL-60 cells (Collins et al., 1978) and murine MPRO (Lawson et al., 1998) and 32D clone 3 (Valtieri et al., 1987) cell lines. More recently, an improved approach was described for deriving neutrophils from murine myeloid progenitors that are conditionally immortalized through controlled expression of HoxB8 (Wang et al., 2006). Expression of HoxB8, a member of the homeobox family of transcription factors, blocks the terminal differentiation of progenitors into monocytes or granulocytes (Krishnaraju et al., 1997; Knoepfler et al., 2001). Neutrophils differentiated *in vitro* from HoxB8-conditional murine myeloid progenitors have been extensively characterized and have shown much promise in replicating the biological functions displayed *ex vivo* by isolated murine neutrophils (McDonald et al., 2011; Weiss et al., 2018; Zehrer et al., 2018; Chu et al., 2019; Saul et al., 2019).

Recent studies have further extended the utilization of HoxB8-conditional progenitors as an experimental tool for *in vivo* studies via their transplantation into mice. Among the wide range of genetic disorders that impact neutrophil development or function, there are more than a dozen different genes whose mutation underlies severe congenital neutropenia as component of the phenotype (Ley et al., 2018). Deficiency of glucose-6-phosphatase-β (G6PC3), an enzyme involved in glucose metabolism, results in neutropenia in humans (Boztug et al., 2009) and mice (Cheung et al., 2007). Utilizing HoxB8-conditional progenitors derived from mice deficient in G6PC3, the neutropenic phenotype was replicated and enabled the investigators to gain mechanistic insight into the role of G6PC3 in neutrophil development (Gautam et al., 2013). In those studies, the HoxB8-conditional progenitor system permitted the exogenous expression of the anti-apoptotic protein Bcl-X_L_ to disentangle the defects in neutrophil progenitor survival from defects in differentiation (Gautam et al., 2013). In another study that employed a mouse model of arthritis, neutrophils derived *in vivo* from HoxB8-conditional progenitors that were adoptively transferred into conditioned mice could be recruited to the inflamed joint (Orosz et al., 2021). In the absence of endogenous host neutrophils, mice that received HoxB8-conditional progenitors exhibited ankle swelling and a clinical score that mirrored that of control mice with intact endogenous neutrophils (Orosz et al., 2021).

There is accumulating evidence that HoxB8-conditional progenitors could be a tool for neutrophil biologists to perform mechanistic and disease model studies *in vivo* on primary-like neutrophils. However, the most significant limitation and drawback to the work described above is that in each study recipient mice needed to be conditioned with irradiation to allow repopulation of the hematopoietic compartment with HoxB8-conditional progenitors. Myeloablative conditioning results in irradiation- or chemical-induced toxicity and inflammation that can significantly impact mouse disease models and confound mechanistic studies (Xun et al., 1994; Duran-Struuck and Dysko, 2009). Orosz and colleagues recently demonstrated the absolute requirement for lethally irradiating recipient mice to achieve engraftment of their HoxB8-conditional progenitors (Orosz et al., 2021). Here, we describe the novel discovery of an independently derived HoxB8-conditional progenitor cell line that is able to robustly engraft in naïve recipient mice, in the absence of conditioning the hematopoietic niche. Our results also indicate that neutrophils generated *in vivo* from transplanted HoxB8-conditional progenitors are functionally equivalent to endogenous primary neutrophils, in terms of their trafficking and effector activities.

## Materials and Methods

### HoxB8 conditionally-immortalized progenitors

As described previously (Cohen et al., 2021), a lentivirus-based construct with a tamoxifen-inducible *Hoxb8* oncogene (Salmanidis et al., 2013) (a gift from Paul Ekert, Murdoch Children’s Research Institute) with either a puromycin or hygromycin resistance cassette was used to stably express HoxB8 in bone marrow hematopoietic stem and progenitor cells (HSPCs) isolated from C57BL/6J mice. HSPCs were cultured with recombinant murine stem cell factor (SCF) and interleukin-3 (IL-3) for two days, exactly as described (Wang et al., 2006), and then subjected to spin-infection (2000 RPM, 60 min, 30°C) with lentivirus in a 24-well plate. Cell culture media consisted of Opti-MEM containing GlutaMax (ThermoFisher), 30 µM beta-mercaptoethanol (Sigma-Aldrich), 10% fetal bovine serum (Gemini Bioproducts), penicillin/streptomycin (Gibco), and non-essential amino acids (Gibco). To establish conditionally-immortalized progenitor lines, cells were cultured in media supplemented with 50 ng/mL SCF (BioLegend), 100 nM Z-4-hydroxytamoxifen (Tocris), and either puromycin (Sigma-Aldrich) or hygromycin (Sigma-Aldrich) for up to two weeks. In some cases, we used conditioned media from CHO cells that secrete recombinant murine SCF (a gift from Patrice Dubreuil, Centre de Recherche en Cancérologie de Marseille).

### Genetic disruption using CRISPR/Cas9

To generate gene knockout cell lines, we used pLentiCRISPR v2 vector, a gift from Feng Zhang (Addgene plasmid #52961). The following 20-nucleotide single-guide RNA (sgRNA) targeting sequences were inserted into pLentiCRSIPR v2: CGGAAGCGAGGTGCAGACCG (*Itga4*), AGTGACATAGAGAATCCCAG (*Itgb1*), AATGTCATCGCGGCGCTCAC (*Cxcr2*); *Tln1* and *Itgb2* targeting have been previously described (Wilson et al., 2017). Lentivirus was produced using a HEK293T Lenti-X cell line (Takara Bio). HoxB8-conditional progenitors were transduced with lentivirus in the presence of 5 µg/mL polybrene (Sigma-Aldrich) in a 24-well plate subjected to centrifugation at 800 x *g* for 60 min.

### Animals and HoxB8-conditional progenitor transplantation

These studies were approved by the Lifespan Animal Welfare Committee (Approval # 5017-19, Office of Laboratory Animal Welfare Assurance #A3922-01) and were conducted in accordance with Public Health Service guidelines for animal care and use. Experiments used 8-14-week old C57BL/6 or CD45.1 (B6.SJL-PtprcaPepcb/BoyJ) mice (The Jackson Laboratory). Water and standard chow were available *ad libitum*. For cell transplantation, CD45.1 mice were anesthetized with 3% isoflurane and received 1 × 10^8^ (unless otherwise indicated) HoxB8-conditional progenitors via retro-orbital i.v. injection.

### Antibodies

Unless otherwise specified, the following antibodies were from BioLegend: anti-Ly6G (1A8), anti-CD45.1 (A20), anti-CD45.2 (104), anti-cKit (ACK2), anti-CXCR4 (FAB21651, R&D Systems), anti-CD11a (M17/4), anti-CD11b (M1/70), anti-CD18 (M18/2), anti-CD29 (HMB1-1), anti-CD49d (R1-2), anti-CD49e (5H10-27), anti-β7 integrin (FIB27), anti-CD44 (IM7), anti-L-selectin (MEL-14), anti-CD16/32 (93), anti-Ly6C (HK1.4), anti-CXCR2 (SA044G4), anti-CD101 (Moushi101, Invitrogen), anti-FcγRIV (9E9), anti-CD162 2PH1, BD Biosciences).

### Analyses of blood and bone marrow

At the indicated time after transplantation of HoxB8-conditional progenitors, blood samples were collected by saphenous venipuncture in K3-EDTA tubes (Sarstedt). Blood samples were subjected to erythrocyte lysis with ammonium chloride (BioLegend). For harvesting bone marrow, mice were euthanized and then femurs and tibias were flushed with ice cold PBS containing 5 mM EDTA and 0.5% BSA (PEB buffer) through 70-µm nylon filters. Samples in PEB buffer were labeled with antibodies as indicated, washed extensively with PEB buffer, and then analyzed by flow cytometry using a MACSQuant Analyzer 10. Data analysis was performed using FlowJo software (BD Biosciences).

### Mouse model of thioglycollate-induced peritonitis

On day 7 post-transplant, an initial blood sample was collected from CD45.1 recipient mice. Mice then received intraperitoneal injection of 1 mL 4% thioglycollate broth (Sigma-Aldrich). Approximately 30 min prior to the end point, a “post-thio” blood sample was collected. At 6 hr after injection of thioglycollate, mice were euthanized and the peritoneal lavage was collected by flushing the peritoneal cavity with 5 mL ice cold PEB buffer. Blood and peritoneal lavage samples were labeled with antibodies against CD45.1, CD45.2, Ly6G, CD18 and/or CXCR2 as indicated, and then analyzed by flow cytometry as described above.

### Phagocytosis assay

Bone marrow was harvested as described above and split into samples of approximately 2.5 × 10^5^ cells each in Hanks’ balanced salt solution containing Ca^2+^/Mg^2+^ (HBSS) and 10% FBS. Some samples were pretreated with 10 µg/mL cytochalasin D prior to adding 10 µg/mL *S. aureus* bioparticles conjugated with a pHrodo-Green fluorescent probe (ThermoFisher Scientific). After allowing phagocytosis by incubating samples at 37°C for 60 min, cells were washed twice with ice cold PEB buffer. Samples were labeled with antibodies against CD45.2 and Ly6G before analysis by flow cytometry as described above.

### *S. aureus* intracellular killing assay

A GFP-expressing strain of *S. aureus*, USA300-sGFP (a gift from Alexander Horswill, University of Colorado Denver), was grown overnight in tryptic soy broth, then diluted 1:10 in fresh broth and grown to log phase. After extensive washing in PBS, the colony forming unit (CFU) density of USA300-sGFP was quantified by measuring its optical density at 600 nm (OD600) on a spectrophotometer. Anti-coagulated blood samples from recipient mice were subjected to lysis of erythrocytes with ammonium chloride (BioLegend) and then labeled at room temperature with antibodies against Ly6G and CD45.2. After washing, samples were resuspended in HBSS containing 10% FBS. Vehicle control or USA300-sGFP (6 × 10^5^ CFUs/sample) was added to samples and they were incubated at 37°C for 15 min to allow *S. aureus* uptake. To remove remaining extracellular *S. aureus*, samples were washed in the presence of 10 mg/mL lysostaphin, pelleted, and resuspended in HBSS containing 10% FBS. A portion of the sample was transferred to ice for analysis by flow cytometry as the initial 0 min time point. The remaining sample was incubated at 37°C, then transferred to ice for analysis by flow cytometry as described above.

### Respiratory burst assay

Anti-coagulated blood samples from recipient mice were diluted 10-fold in HBSS containing Ca^2+^/Mg^2+^. Samples were then loaded for 15 min with 2.5 µg/mL dihydrorhodamine 123 (DHR123; Cayman Chemical), a probe that fluoresces upon exposure to reactive oxygen intermediates. After DHR123 loading, diphenyleneiodonium (DPI; Sigma Aldrich) was added to the appropriate samples at a concentration of 10 µM just prior to stimulation of samples with either heat-killed *S. aureus* (HKSA) or 100 ng/mL phorbol 12-myristate 13-acetate (PMA; Sigma-Aldrich). Samples were stimulated for either 15 min (PMA) or 30 min (HKSA). At the end of stimulation, samples were centrifuged to pellet cells, subjected to red blood cell lysis as described above, and then stained on ice with antibodies against CD45.2, CD45.1, and Ly6G. Samples were analyzed by flow cytometry as described above.

### Statistical analyses

All statistical analyses were performed using Prism 8 (GraphPad). Data throughout are presented as the group mean and standard deviation. One-way or two-way analysis of variance (ANOVA) was used to compare the differences between group means and post-hoc analysis was performed using a Bonferroni multiple comparison test.

## Results

### Engraftment of HoxB8-conditional progenitors

Since the approach for conditionally immortalizing murine myeloid progenitors by exogenous expression of HoxB8 was initially described (Wang et al., 2006), others have shown that such progenitor cell lines can be transplanted into irradiated recipient mice and subsequently generate mature, terminally differentiated myeloid leukocytes in the *in vivo* environment (Gautam et al., 2013; Redecke et al., 2013; Orosz et al., 2021). Recently, we reported that a HoxB8-conditional progenitor cell line that we derived is able to produce substantial numbers of donor graft-derived neutrophils in mice that were not subjected to any conditioning of the hematopoietic niche (Cohen et al., 2021). To understand if the engraftment of HoxB8-conditional progenitors in naïve, unconditioned mice was a unique feature of this particular progenitor cell line, we compared its engraftment to other lines of HoxB8-conditional myeloid progenitors using congenic CD45.1 recipient mice. Each of the three cell lines tested were independently generated by lentiviral delivery of a tamoxifen-inducible *Hoxb8* transgene to hematopoietic stem and progenitor cells (HSPCs) isolated from the bone marrow of C57BL/6J mice (CD45.2^+^). To determine whether three distinct HoxB8-conditional progenitor lines would home to and take up residence in the bone marrow, we transplanted them separately in CD45.1 mice that had not received any prior conditioning regimen. For two of the independently-established progenitor lines, line 2 and line 3, we observed very few donor-derived cells present in the recipient mouse bone marrow at 4 days after transplant (Figure 1A). This was in contrast to the robust engraftment of progenitor line 1, as measured by the fraction of CD45.2^+^ cells in bone marrow (among all CD45^+^ cells) that were derived from transplanted donor progenitors (Figure 1A). These results indicate that HoxB8-conditional progenitor line 1 has a unique capacity for hematopoietic-intact engraftment.

**Figure 1.**
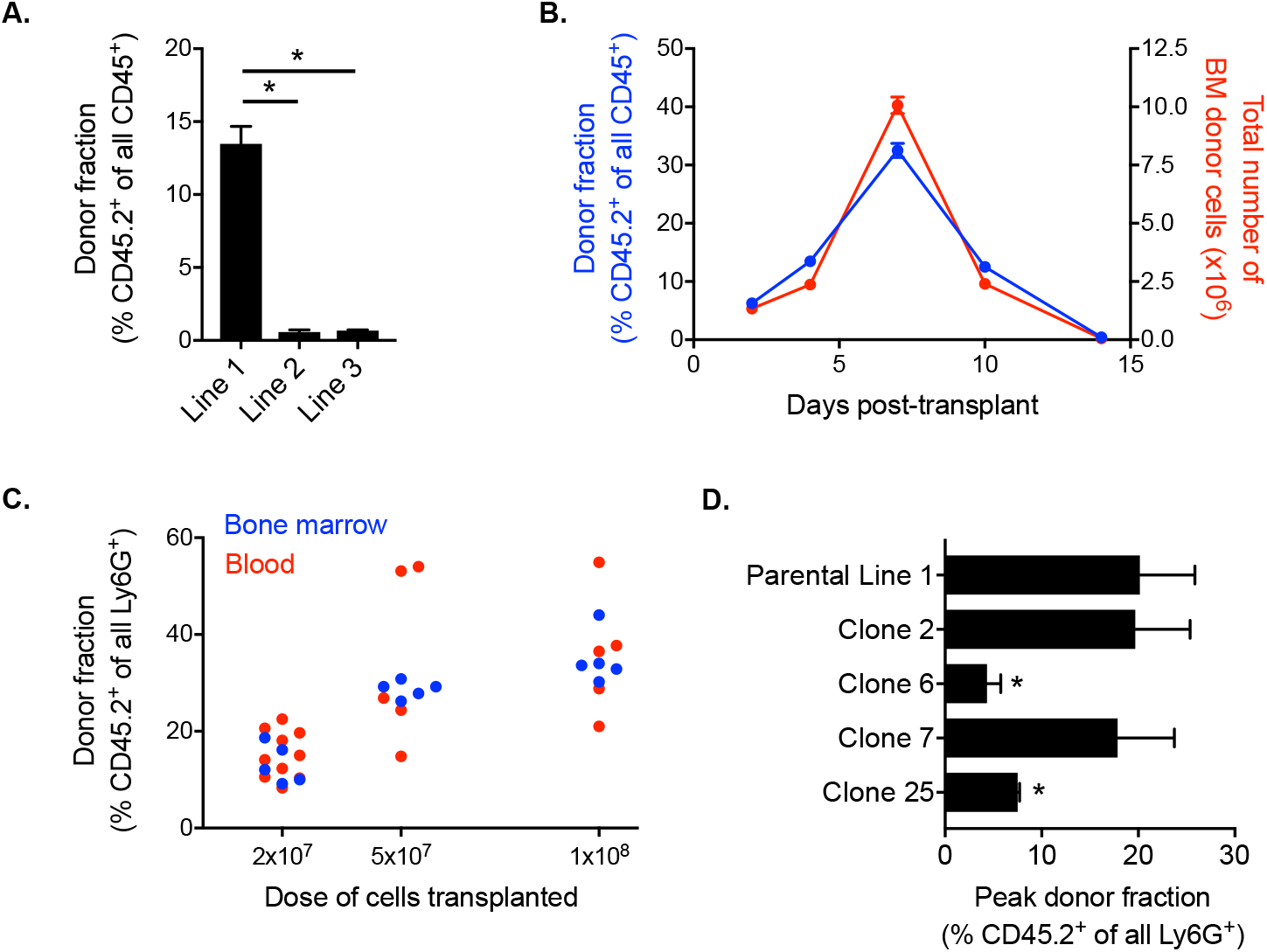
Quantification of HoxB8-conditional progenitor engraftment in unconditioned mice. **(A)** CD45.1 mice received transplantation of 1 × 10^8^ progenitors of one of the indicated HoxB8-conditional progenitor cell lines. The fraction of all CD45^+^ bone marrow cells that were derived from transplanted HoxB8-conditional progenitors (CD45.2^+^) in bone marrow of recipient mice at five days after transplantation. **(B)** CD45.1 mice received transplantation of 1 × 10^8^ HoxB8-conditional progenitors (line 1) to perform a time course of both the fraction (blue, left y-axis) and absolute number (red, right y-axis) of donor-derived cells in the bone marrow. **(C)** The donor-derived (CD45.2^+^) fraction of all mature Ly6G^high^ neutrophils in the bone marrow CD45.1 mice that received varying doses of HoxB8-conditional progenitors, measured at seven days post-transplant. **(D)** Evaluation of the maximum fraction of donor-derived (CD45.2^+^) neutrophils in the peripheral blood of CD45.1 mice that received transplant of 3 × 10^7^ cells of either the parental HoxB8-conditional progenitor cell line 1 or one of the single-cell clones derived from that cell line.

To further characterize the engraftment of HoxB8-conditional progenitor cell line 1, we used CD45.1 recipient mice and performed a kinetic analysis of donor-derived CD45.2^+^ cells in the bone marrow. In this experiment, CD45.1 received 1 × 10^8^ HoxB8-conditional progenitors on day 0. The fraction of all bone marrow CD45^+^ cells that are derived from transplanted donor cells peaks around day 7 after transplant and are nearly undetectable by day 14 (Figure 1B), following kinetics similar to those observed by others in irradiated recipient mice (Orosz et al., 2021). The absolute number of CD45.2^+^ donor-derived cells in the bone marrow follows a similar trajectory (Figure 1B), suggesting that a relatively small fraction of transplanted HoxB8-conditional progenitors initially engraft and that it is their proliferation within the hematopoietic niche that drives the substantial donor chimerism that develops over the first week following transplant. Furthermore, we observed a dose-dependence between the number of transplanted progenitors and the fraction of neutrophils in the bone marrow and peripheral blood that were donor-derived on day 7 after transplant (Figure 1C). At longer time points of 45 days and 6 months, we did not detect any donor-derived cells in the bone marrow and there was not a detectable long-term impact on the host stem/progenitor cell composition of the bone marrow (Figure S1).

The derivation of HoxB8-conditional progenitors from bone marrow HSPCs is likely to produce a heterogeneous mix of conditionally-immortalized myeloid progenitor states. To further understand whether the disparate engraftment among the three HoxB8-immortalized cell lines was due to the presence of a subpopulation with an intrinsic ability to engraft in the bone marrow niche, we generated single-cell clones of progenitor line 1 and then analyzed the engraftment of four distinct clonal progenitor lines that were transplanted into naïve CD45.1 recipient mice. We analyzed peripheral blood longitudinally to assess the peak fraction of donor-derived neutrophils in the periphery among the different clonal lines tested, as there was some variability between progenitor clones in their kinetics of engraftment and differentiation. In addition, we utilized a transplantation dose of 3 × 10^7^ HoxB8-conditional progenitors in order to better resolve potential differences between progenitor line 1 and the clonal progenitor lines. Across four different single-cell clones, we observed a range in the peak fraction of peripheral blood neutrophils that were derived from transplanted donor progenitors (Figure 1D). Clones 6 and 25 exhibited a reduced peak frequency of donor-derived neutrophils in the periphery compared to the parental line 1, whereas the donor fraction for clones 2 and 7 was comparable to that of the parental progenitor line 1 (Figure 1D). These data suggest the presence of a subset of highly-engraftable progenitors within the parental HoxB8-conditional progenitor line 1. As these analyses involved serial sampling of the peripheral blood, we accrued a substantial number of data points to be able to assess whether neutrophils generated from transplanted donor progenitors displace those generated via endogenous host granulopoiesis. There was no detectable relationship between the host-derived neutrophil count in the blood and the fraction of all neutrophils that were derived from the donor (Figure S1).

To gain insight into potential cell surface receptors that may regulate engraftment or define a subset of highly-engrafting neutrophil progenitors, we performed flow cytometry analyses of HoxB8-conditional progenitor lines and single-cell clones to evaluate expression of cKit; C-X-C chemokine receptor 4 (CXCR4); integrin subunits CD11a, CD11b, and CD29; CD44, L-selectin, CD16/32, CD101, Ly6C, and Ly6G. Most of these receptors and markers exhibited homogeneous expression levels within each line (Figure S2) and many had similar expression across the different progenitor cell lines analyzed (Figure 2). All progenitor lines lacked expression of Ly6G and CD101 (Figure S2). Notable differences between progenitor lines include higher cKit expression for the poorly engrafting line 2 and disparate expression levels of several receptors/markers (CD11a, CD29) on progenitor line 3 (Figure 2). There was a prominent Ly6C^high^ population within line 1, line 3, and all of the clonal progenitor lines derived from progenitor line 1 (Figure 2). Compared to line 1, line 2 had a reduced fraction of Ly6C^high^ progenitors while line 3 exhibited a near 100% fraction of the Ly6C^high^ population (Figure 2). However, the frequency of the Ly6C^high^ subpopulation exhibited no apparent correlation with the intrinsic engraftment capacity of the respective HoxB8-conditional progenitor cell lines. Together, these data suggest that conditional HoxB8 expression in myeloid progenitors may vary in the state or identity of the progenitor that is immortalized.

**Figure 2.**
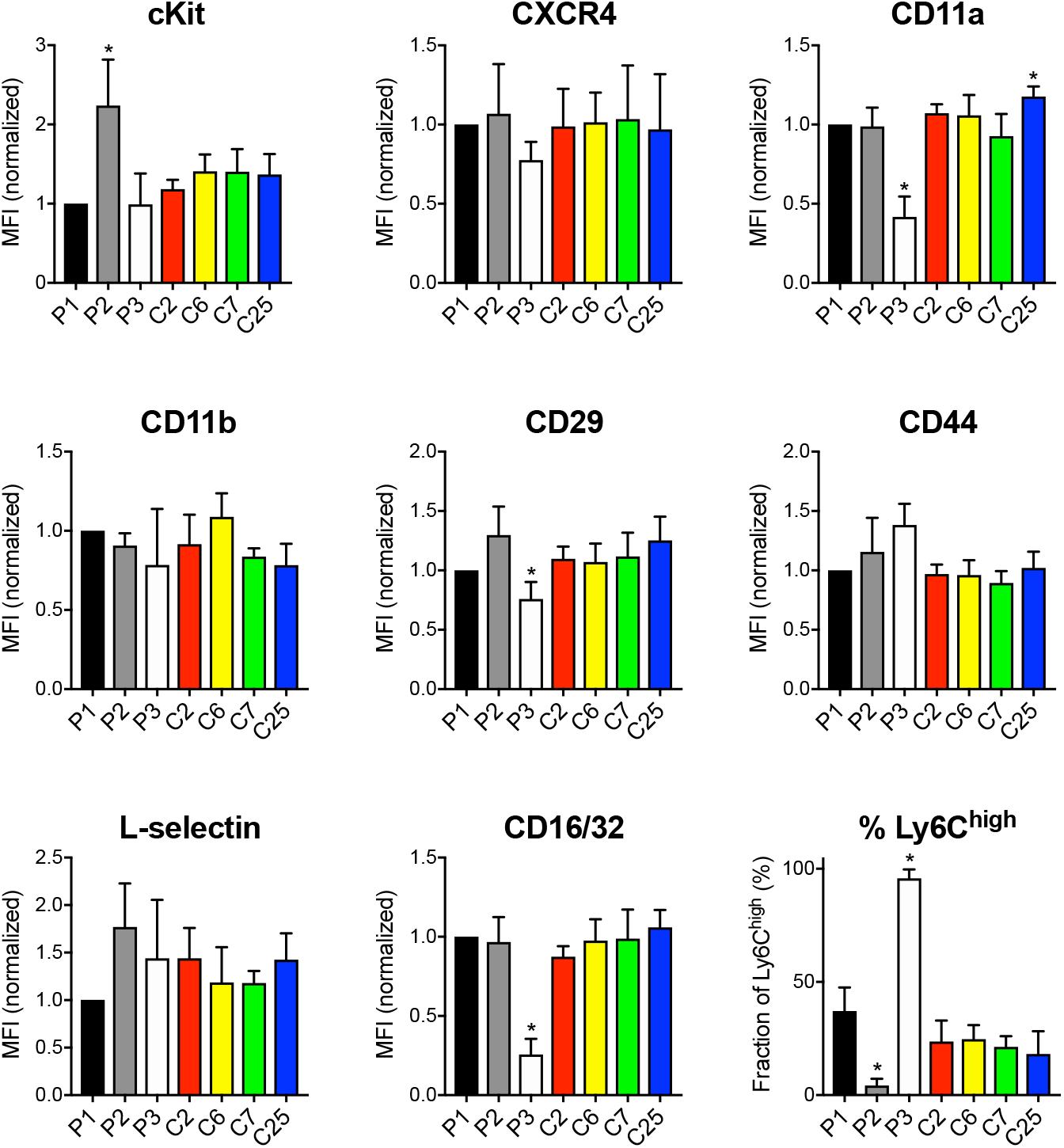
Receptor and surface marker expression on HoxB8-conditional progenitors. Flow cytometry analyses was performed to quantify the cell surface expression of the indicated receptors and markers on the three main HoxB8-conditional progenitor cell lines; line 1 (P1), line 2 (P2), line 3 (P3); and the P1-derived single-cell clonal lines designated clone 2 (C2), clone 6 (C6), clone 7 (C7), and clone 25 (C25).

### Mechanisms of HoxB8-conditional progenitor engraftment

Prior to taking up residence in the hematopoietic niche, transplanted HSPCs must first traffic from the circulation to the bone marrow. Broadly, HSCs and HPCs transplanted into conditioned recipients undergo engraftment that depends on the integrin VLA-4 (Papayannopoulou et al., 1995; Potocnik et al., 2000; Qian et al., 2007; Rettig et al., 2012). To determine whether VLA-4 plays a role in the engraftment of HoxB8-conditional progenitors, we used CRISPR/Cas9 to generate progenitor cell lines deficient in either *Itga4* or *Itgb1* that encode the α4 and β1 subunits of VLA-4, respectively (Figure S3). Congenic CD45.1 mice received a 1:1 mix of wild-type and either *Itga4*^*-/-*^ or *Itgb1*^*-/-*^ HoxB8-conditional progenitors. At five days post-transplant, we measured the relative fraction of wild-type and knockout CD45.2^+^ donor-derived cells in the bone marrow (Figure 3A). Interestingly, in the respective recipient mice, the abundance of *Itga4*^*-/-*^ donor cells was similar to that of wild-type donor cells (Figure 3A). In contrast, *Itgb1*^*-/-*^ HoxB8-conditional progenitors exhibited a level of engraftment that was significantly less than that of wild-type (Figure 3A). These data indicate that VLA-4 is not essential for the engraftment of HoxB8-conditional progenitors, but that another β1 integrin subtype plays a role in this process. Flow cytometry analyses of the *Itga4*^-/-^ progenitor cell line indicated that CD29 (β1 integrin subunit) remains expressed at the cell surface (Figure S3), indicating that other β1 integrins are expressed. Wild-type, but not *Itgb1*^*-/-*^, progenitors expressed CD49e and CD49f that pair with CD29 to form α5β1 and α6β1 integrins, respectively (Figure S3). Wild-type, but not *Itga4*^*-/-*^, progenitors also expressed α4β7 integrins (Figure S3).

**Figure 3.**
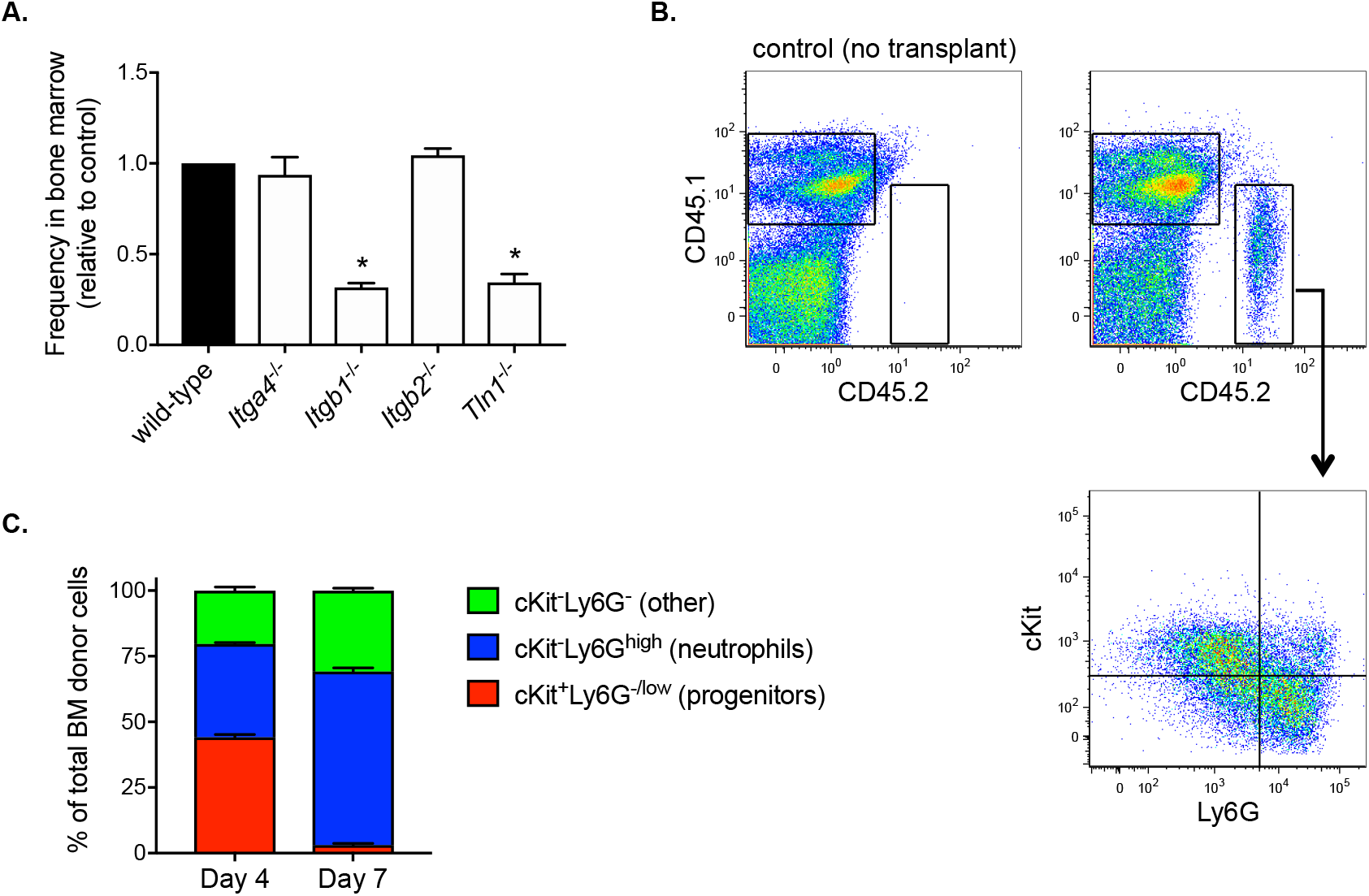
HoxB8-conditional progenitor engraftment and *in vivo* differentiation. **(A)** Five days after CD45.1 mice received a 1:1 mix of wild-type and the indicated gene-deficient HoxB8-conditional progenitors, the relative frequency of wild-type and gene knockout cells among all donor-derived cells in the bone marrow was measured by flow cytometry. Data are presented as the relative frequency of each gene knockout type relative to its wild-type counter part, and normalized to the wild-type control (set to equal 1.0). **(B)** Example flow cytometry dotplots showing analyses of whole bone marrow from CD45.1 mice that received transplant of CD45.2^+^ HoxB8-conditional progenitors. Samples were labeled to determine expression of CD45.1, CD45.2, cKit and Ly6G. Gating for analyzing only donor-derived cells (top) and the subpopulations of donor-derived cells (bottom) in various states of maturity from progenitors/immature neutrophils (cKit^+^Ly6G^-/low^) towards terminal neutrophils (cKit^-^Ly6G^high^). **(C)** Characterization and quantification of the *in vivo* differentiation of engrafted HoxB8-conditional progenitors at day 4 and day 7 after transplantation.

Extravasation of some leukocyte subsets from blood vessels is dependent on β2 integrins and a recent study suggests that β2 integrins contribute to transplanted HSPC homing (Mei et al., 2020). In contrast to our observations of the reduced engraftment of β1 integrin-deficient HoxB8-conditional progenitors, *Itgb2*^*-/-*^ donor progenitors (Figure S3) were present in the bone marrow at similar frequencies as wild-type when transplanted at a 1:1 ratio (Figure 3A). Talin binds to the cytosolic tail of all integrin β chains and is required for integrin activation (Kim et al., 2011). Similar to β1 integrin-deficient HoxB8-conditional progenitors, talin knockout (*Tln1*^-/-^) progenitors exhibited significantly reduced engraftment compared to wild-type control (Figure 3A).

If transplanted donor HoxB8-conditional progenitors truly engraft and take up residence in the bone marrow niche, even if that residence is transient, one would expect to observe progenitor cell proliferation and progressive differentiation towards terminal leukocyte states. We have previously shown that cells in the bone marrow derived from transplanted HoxB8-conditional progenitors that were loaded with a cell proliferation dye exhibit a greater than 20-fold dilution of the dye within 3 days, indicating *in vivo* proliferation of donor progenitors prior to their differentiation (Cohen et al., 2021). Analyses of all donor-derived cells in the bone marrow at days 4 and 7 after transplant indicate that their composition shifts from being split between cKit^+^Ly6G^low^ progenitors and cKit^-^Ly6G^high^ neutrophils at day 4, to predominantly cKit^-^ Ly6G^high^ neutrophils at day 7 (Figure 3B and 3C). Taken together, these data suggest that HoxB8-conditional progenitors home to the bone marrow and engraft in the hematopoietic niche, where they transiently proliferate and generate large numbers of mature neutrophils for up to two weeks.

### Trafficking of neutrophils derived from transplanted HoxB8-conditional progenitors

Several studies have demonstrated that HoxB8-conditional progenitors can produce mature neutrophils following their transplantation into irradiated mice (Gautam et al., 2013; Redecke et al., 2013; Orosz et al., 2021). To further probe the dynamics of neutrophils generated *in vivo* from transplanted HoxB8-conditional progenitors, we first performed experiments using a model of sterile peritonitis induced by intraperitoneal injection of thiogylcollate. First, CD45.1 mice received transplantation of wild-type HoxB8-conditional progenitors and given sufficient time to produce robust chimerism in the blood. In response to peritonitis, we observed, as indicated by the similar levels of donor chimerism in the blood (before and after thioglycollate injection) and in the inflamed peritoneal cavity (Figure 4A). These data indicate that donor-derived neutrophils are recruited to a site of inflammation to an equal extent as endogenous host neutrophils.

**Figure 4.**
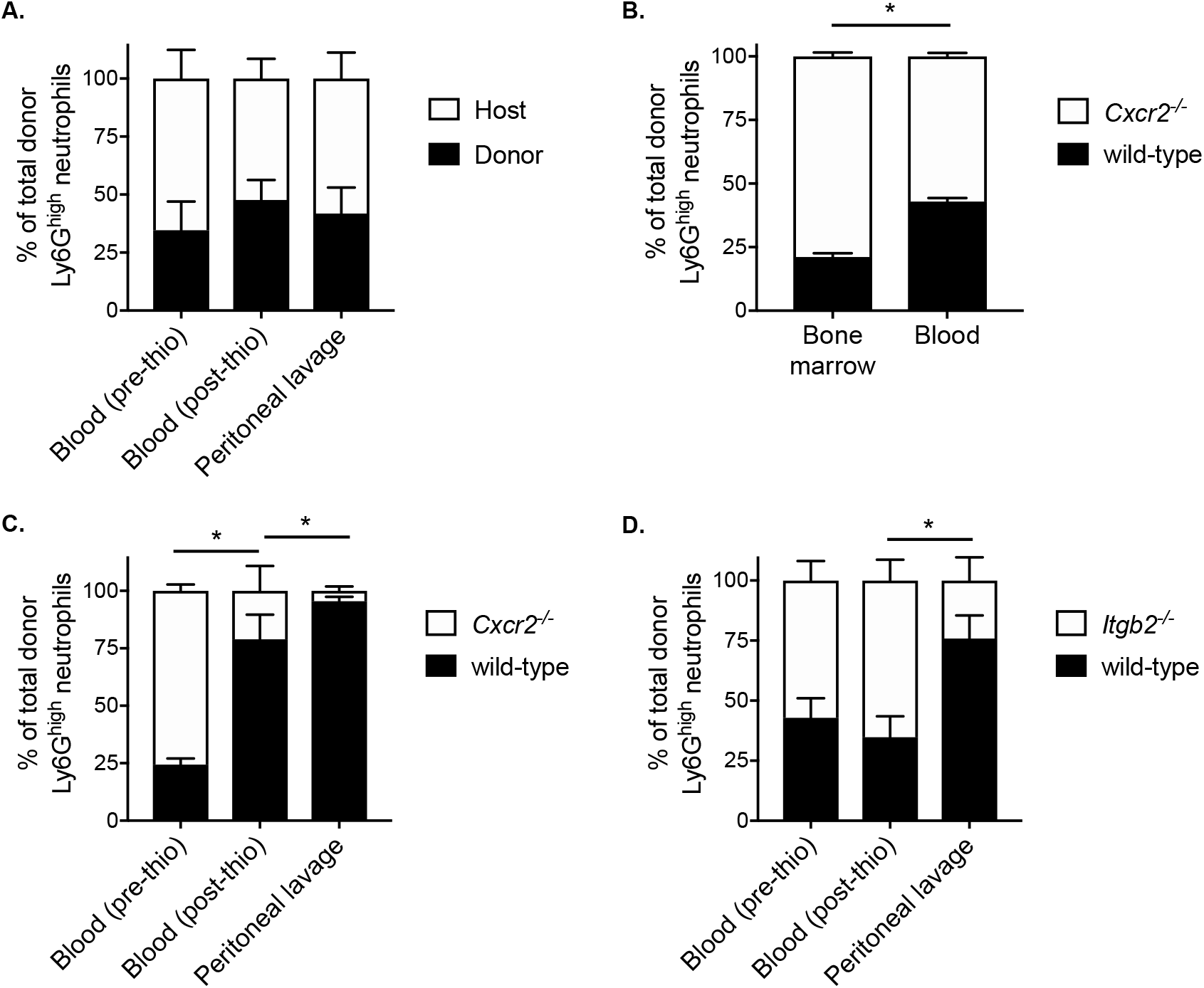
Mechanisms underlying mobilization and recruitment of donor-derived neutrophils. Wild-type HoxB8-conditional progenitors, alone **(A)** or mixed with CXCR2-deficient **(B, C)** or β2 integrin-deficient HoxB8-conditional progenitors **(D)**, were transplanted into CD45.1 recipient mice. **(A, C, D)** Mice were subjected to sterile peritonitis induced by thioglycollate. Blood samples (before and after thioglycollate injection) and peritoneal lavage were analyzed for the frequency of donor-derived wild-type and gene knockout neutrophils, as indicated. **(B)** In mice not subjected to any inflammatory stimulus, bone marrow and blood samples were collected at day 8 post-transplant and analyzed for the frequency of donor-derived wild-type and CXCR2-deficient neutrophils.

Neutrophils mature in the bone marrow and their subsequent mobilization to the peripheral circulation is regulated by C-X-C chemokine receptor 2 (CXCR2) (Eash et al., 2010). To further understand whether neutrophils derived *in vivo* from transplanted progenitors are mobilized to the periphery and recruited to a site of inflammation via mechanisms that have been described for primary neutrophils, we performed several types of analyses in mice that had received a mix of wild-type and gene knockout donor progenitors. In these studies, CD45.1 mice received both wild-type and *Cxcr2*^*-/-*^ HoxB8-conditional progenitors and we restricted our analyses to CD45.2^+^Ly6G^high^ donor neutrophils (Figure S4), as nearly 100% of wild-type neutrophils (both host- and donor-derived) express CXCR2 (Figure S5). At day 8 post-transplant, donor neutrophils of both genotypes were present in the bone marrow and peripheral blood of recipient mice. However, there was a significant reduction in the relative fraction of those donor neutrophils that were *Cxcr2*^*-/-*^ in the blood compared to the bone marrow (Figure 4B). These data indicate that homeostatic egress of mature donor-derived neutrophils to the periphery is regulated by CXCR2 in a similar manner to that of primary endogenous neutrophils (Eash et al., 2010).

To evaluate the role of CXCR2 in the mobilization of donor neutrophils in response to inflammatory stimuli, we analyzed the cellular composition of peripheral blood before and after mice received intraperitoneal thioglycollate to induce a sterile peritonitis. Similar to the analyses of homeostatic neutrophil egress to the periphery, the CD45.1 mice first received a transplant of a mix of CD45.2-expressing wild-type and *Cxcr2*^*-/-*^ HoxB8-conditional progenitors. The mix of donor progenitors for transplantation was composed of approximately one-third wild-type and two-thirds *Cxcr2*^*-/-*^ progenitors, as we wanted to be able to detect changes in donor neutrophil fraction with the expectation that CXCR2 is involved both in neutrophil mobilization and recruitment, as is known for primary murine neutrophils (Cacalano et al., 1994; Eash et al., 2010). At 90 minutes after inducing inflammation, there was a significant decrease in the fraction of donor-derived CXCR2-deficient neutrophils in the blood (Figure 4C), indicating a CXCR2-dependent mobilization of donor neutrophils from the bone marrow. There was a further reduction in the fraction of donor-derived *Cxcr2*^*-/-*^ neutrophils relative to wild-type in the peritoneal lavage, relative to the blood (Figure 4C). Taken together, these data confirm that both the stimulated mobilization and recruitment of neutrophils generated *in vivo* from transplanted HoxB8-conditional progenitors occurs via CXCR2-dependent mechanisms.

Endogenous host neutrophils and those derived *in vivo* from transplanted donor HoxB8-conditional progenitors express similar levels of adhesion and signaling receptors involved in regulating neutrophil recruitment, including PSGL-1, L-selectin, CD11a, CD11b, CXCR2, and FcγRIV (Figure S5). These data are consistent with a recent study, with the exception that we observed equal expression of CD11b and FcγRIV on donor and host neutrophils (Figure S5), whereas those receptors were elevated on neutrophils when produced in irradiated recipient mice (Orosz et al., 2021). Two of the major types of β2 integrins expressed by neutrophils, LFA-1 (αLβ2) and Mac-1 (αMβ2), play key roles in neutrophil recruitment by mediating adhesion to inflamed endothelium (Ley et al., 2007). To understand whether donor-derived neutrophils are recruited to sites of inflammation via adhesion mechanisms that have been well described for host-derived primary murine neutrophils, we again used a model of thioglycollate-induced peritonitis in mice that received a 1:1 mix of wild-type and β2 integrin-deficient (*Itgb2*^-/-^) HoxB8-conditional progenitors. While there was a similar proportion of wild-type and *Itgb2*^-/-^ donor neutrophils in the blood both before and after inducing inflammation, we observed a significant decrease in the fraction of β2 integrin-deficient donor-derived neutrophils in the peritoneal lavage (Figure 4D). These data indicate that, like primary host neutrophils, neutrophils derived *in vivo* from transplanted HoxB8-conditional progenitors are recruited to the inflamed peritoneum via β2 integrin-dependent mechanisms.

### Effector functions of donor neutrophils derived *in vivo*

Previous studies have characterized the effector functions of mature neutrophils derived *in vitro* and *in vivo* from HoxB8-conditional progenitors (McDonald et al., 2011; Weiss et al., 2018; Zehrer et al., 2018; Chu et al., 2019; Saul et al., 2019; Orosz et al., 2021). Some of these studies are limited to evaluating only the function of neutrophils derived from HoxB8-conditional progenitors *in vitro* or lack a controlled *in vivo* comparison with endogenous host primary neutrophils. Since the unique line of HoxB8-conditional progenitors that we describe in this study engrafts in unconditioned mice, we were therefore able to directly assay both host and donor neutrophil functions under internally controlled conditions. Using CD45.1 mice that received CD45.2^+^ donor progenitors, we assayed neutrophil effector functions using blood samples. First, we exposed blood samples to *Staphylococcus aureus* bioparticles conjugated to a pHrodo-Green dye that fluoresces upon internalization into the neutrophil phagolysosome, using a low particle:cell ratio so as to better resolve potential differences between host- and donor-derived neutrophils (Figure S6). We observed that neutrophils derived *in vivo* from transplanted HoxB8-conditional progenitors had a similar capacity for *S. aureus* bioparticle phagocytosis that was inhibited by the actin-disrupting agent cytochalasin D (Figure 5A).

**Figure 5.**
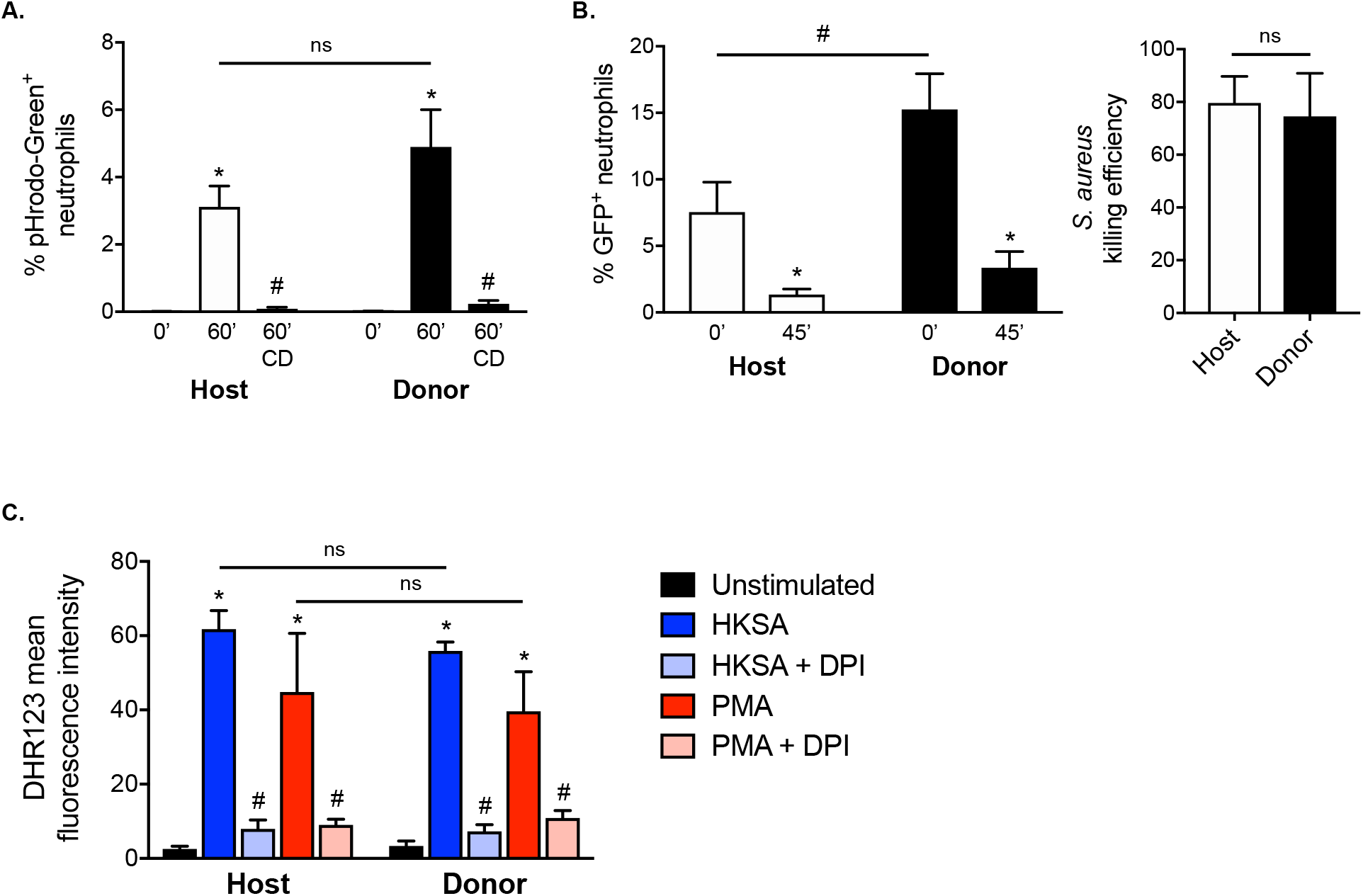
Effector function of neutrophils derived *in vivo* from HoxB8-conditional progenitors. Seven days after CD45.1 mice received transplant of wild-type HoxB8-conditional progenitors, blood samples were subjected to assays of neutrophil function using flow cytometry. CD45.2^+^Ly6G^+^ donor-derived neutrophils were distinguished from CD45.1^+^Ly6G^+^ host-derived neutrophils for comparative analyses. **(A)** Host and donor neutrophil phagocytosis of pHrodo-Green conjugated *S. aureus*, measured after 60 min exposure in the absence or presence of 10 µg/mL cytochalasin D (CD). **(B)** Host and donor neutrophil intracellular killing of *S. aureus*, as measured by the fraction of neutrophils remaining GFP^+^ 45 min after allowing initial phagocytosis of strain USA300-sGFP. The killing efficiency was calculated by comparing the end point fraction of GFP^+^ neutrophils to the starting point (0 min). **(C)** Quantification of stimulated ROS generation by host and donor neutrophils, expressed as the mean fluorescence intensity of DHR123. Blood samples were stimulated with either heat-killed *S. aureus* (HKSA) or 100 ng/mL PMA, in the absence or presence of 10 µM DPI.

To determine whether neutrophils are able to kill *S. aureus* after uptake, we employed an assay based upon the intracellular killing event being shown to result in the bleaching of GFP expressed by a strain of *S. aureus* (Schwartz et al., 2009). Donor-derived blood neutrophils were able to eradicate live *S. aureus* to a similar extent as host-derived neutrophils (Figure 5B). Neutrophil intracellular killing of *S. aureus* occurs in a manner dependent on the NADPH oxidase (NOX2). Both host- and donor-derived neutrophils underwent NOX2-mediated respiratory burst in response to heat-killed *S. aureus* or PMA, and sensitive to the NOX2 inhibitor DPI (Figure 5C). Together, these data indicate that neutrophils generated from HoxB8-conditional progenitors following their transplantation into naïve mice have competent host defense effector functions.

## Discussion

Neutrophils are essential players in the acute inflammatory response. In recent years, investigators have continued to identify novel and diverse roles for neutrophils in many facets of host defense and immune regulation (Kubes, 2018). While neutrophil biologists have long faced experimental limitations and challenges, the emergence of the HoxB8-conditional progenitor system has provided a means for researchers to obtain near-primary neutrophils that are genetically tractable prior to their differentiation. Here, we report the discovery of a unique derivation of a HoxB8-conditional progenitor cell line that is capable of engrafting in naïve mice and results in robust donor neutrophil chimerism that lasts for more than a week.

Hematopoietic stem cell transplantation (HSCT) can be curative for blood cell disorders and malignancies. HSPCs reside in specialized niches within the extravascular space of the bone marrow. For HSCT to be successful, transplanted HSPCs must traffic to these specialized hematopoietic niches and then take up residence through a process called engraftment. It remains a topic of debate whether the engraftment of transplanted HSPCs is limited by the occupancy of specialized niches by host HSPCs (Bhattacharya et al., 2008). Clinically, HSCT recipients undergo irradiation- and/or chemical-induced ablation of the hematopoietic system prior to the injection of HSCs or whole bone marrow. This conditioning of the host promotes donor engraftment through a combination of blocking rejection of the graft by the host immune response and freeing up hematopoietic niche space for donor HSPCs. In the absence of conditioning, while HSC engraftment remains poor even in recipients that are incapable of graft rejection (Muller et al., 2000; Czechowicz et al., 2007; Bhattacharya et al., 2008), the availability of niches for specific subsets of multipotent and lineage-restricted progenitors has not been extensively investigated. However, our derivation of a HoxB8-conditional progenitor cell line that efficiently engrafts in unconditioned mice appears to be a novel discovery, as we found that other independently generated HoxB8-conditional progenitor lines did not engraft and others have demonstrated the absolute requirement for irradiating recipient mice in order to achieve engraftment and donor neutrophil chimerism (Orosz et al., 2021).

Similar to the investigation of HSPCs and conditioning the hematopoietic niche, the mechanisms of HSC homing and engraftment are more clearly defined than those of progenitor subsets. The integrin VLA-4 (α4β1) is essential for the homing of transplanted HSCs to the bone marrow (Papayannopoulou et al., 1995; Potocnik et al., 2000; Qian et al., 2007; Rettig et al., 2012). We were therefore surprised to observe that *Itga4*^-/-^ HoxB8-conditional progenitors lacking VLA-4 expression engrafted and proliferated in the bone marrow of recipient mice as well as wild-type progenitors. By contrast, β1 integrin-deficient HoxB8-conditional progenitors have impaired homing and/or engraftment, suggesting that a β1 integrin subtype other than VLA-4 plays a role in this process. Furthermore, since talin knockout HoxB8-conditional progenitors were also reduced in their bone marrow frequency relative to wild-type, it would suggest a homing mechanism that involves activated β1 integrins, similar to that observed for β2 integrins and neutrophil trafficking (Lefort et al., 2012). Further investigation of the mechanisms of engraftment for HoxB8-conditional progenitors will also provide insight as to how this cell line is uniquely able to engraft in the absence of myeloablation.

In recent studies, single-cell technologies have enabled the identification of the various neutrophil precursor states that define the maturation process following lineage commitment (Evrard et al., 2018; Zhu et al., 2018; Kwok et al., 2020). Our analyses of cell surface receptors and markers reveal that the progenitor identity that is conditionally immortalized through enforced HoxB8 expression appears to vary between several independent derivations, despite following the same protocol. HoxB8-conditional progenitor line 1 that we describe in this study may be best characterized as a granulocyte macrophage progenitor (GMP), given the bifurcated expression of Ly6C by the parental progenitor line 1 and single-cell clones derived from it, a pattern that exists within GMPs and may be related to lineage commitment decisions (Kwok et al., 2020). Comparing the timing of *in vivo* differentiation in our studies and those recently reported for neutrophil-committed preNeus (Evrard et al., 2018), the HoxB8-conditional progenitor lines we characterize are unlikely to be in the preNeu stage while HoxB8 is expressed since we did not observe a substantial shift towards cKit^-^Ly6G^high^ donor cells in the bone marrow until day 4 and later. Future systematic analyses are necessary to understand the divergent outcomes of HoxB8-dependent progenitor immortalization with respect to their developmental identity and eventual engraftment potential as an established cell line.

Our studies confirm that neutrophils derived *in vivo* from donor progenitors are biologically comparable to endogenous neutrophils, building upon the results of others that have utilized transplantation of HoxB8-conditional progenitor into mice (Gautam et al., 2013; Orosz et al., 2021). In order to show this more definitively, the experiments described here are, to our knowledge, the first to extensively probe the molecular mechanisms of trafficking and function of neutrophils generated *in vivo* from transplanted HoxB8-conditional progenitors. From CXCR2-dependent mobilization from the bone marrow to neutrophil recruitment mediated by CXCR2 and β2 integrins, we demonstrate that donor-derived neutrophils are functionally equivalent to endogenous host neutrophils. We found that HoxB8-conditional progenitor differentiation that occurs *in vivo* in the microenvironment of the bone marrow allows donor neutrophils to reach their full functional competence. This contrasts with certain neutrophil functions observed with *in vitro* differentiation of HoxB8-conditional progenitors, under conditions that are unlikely to fully recapitulate the environment in which neutrophil maturation occurs in the bone marrow. For example, NOX2-dependent stimulated generation of ROS is more than an order of magnitude lower for *in vitro*-differentiated neutrophils compared to primary neutrophils (Saul et al., 2019). Our results show that ROS generation in response to either *S. aureus* bioparticles or PMA is no different between endogenous host neutrophils and those derived *in vivo* from transplanted HoxB8-conditional progenitors. In addition, intracellular eradication of *S. aureus* occurs as well in donor-derived neutrophils as in host-derived neutrophils. Taken together, our results strongly suggest that mature neutrophils that are generated in mice from adoptively transplanted HoxB8-conditional progenitors are biologically equivalent to primary neutrophils.

Multidrug-resistant (MDR) bacterial and fungal infections are an increasingly urgent clinical challenge, especially for opportunistic pathogens that cause severe disease in the immunocompromised. Due to a number of factors, the pipeline for new antibiotics has seen a drastic reduction in productivity (Lewis, 2013). Immunomodulatory and cell therapeutic strategies are being pursued as a complementary approach or alternative to antibiotics (Hancock et al., 2012; Kumaresan et al., 2014). While granulocyte transfusion therapy has experienced a long and detoured path in clinical implementation (Marfin and Price, 2015), it may be worth revisiting its conceptual basis as new approaches are developed for generating neutrophils (both *ex vivo* and *in vivo*). We posit that conditionally-immortalized neutrophil progenitors could be an effective cellular therapeutic in the context of neutropenia or a dysregulated inflammatory response. Given their genetic tractability, murine HoxB8-conditional progenitors represent an innovative system for investigating means to enhance neutrophil effector functions. In this way, an off-the-shelf product that transiently generates neutrophils with superior capacity to, for example, overcome evasive intracellular survival of MDR *S. aureus* (Greenlee-Wacker et al., 2015) may be an option alongside antibiotics of last resort.

In conclusion, our studies describe a new HoxB8-conditional progenitor cell line that is uniquely capable of transient engraftment in the unconditioned murine host in a manner that depends on β1 integrins, but not VLA-4. Within the bone marrow, transplanted HoxB8-conditional progenitors proliferate and differentiate into mature neutrophils that are mechanistically and functionally the equivalent of primary neutrophils. These results further confirm the utility of the HoxB8 progenitor system for experimental neutrophil biology and suggest future translational avenues.

## Acknowledgments

We thank Dr. Paul Ekert and Dr. Alexander Horswill for kindly providing reagents/resources.

## Funding

This research was supported by awards from the National Institutes of Health (GM124911 to C.T.L.; GM119943 and CA218079 to P.M.D.; T32GM065085 supported J.T.C.; and T32HL134625 supported B.M.N.), Brown Physicians, Inc. (C.T.L.), and the Armand D. Versaci Research Scholarship in Surgical Sciences (M.D.).

## Author Contributions

J.T.C.: Developing cell lines, designing research studies, conducting experiments, analyzing data.

M.D.: Developing cell lines, designing research studies, conducting experiments, analyzing data.

K.D.H.: Designing research studies, conducting experiments, analyzing data.

B.M.N.: Designing research studies, conducting experiments, analyzing data.

R.J.: Designing research studies, conducting experiments.

Z.S.W.: Developing cell lines.

A.C.: Designing research studies, analyzing data.

P.M.D.: Designing research studies.

C.T.L.: Supervising the project, designing research studies, conducting experiments, analyzing data, writing the manuscript.

## Competing Interests

The authors have declared that no competing interests exist.

---

**Figure S1.**
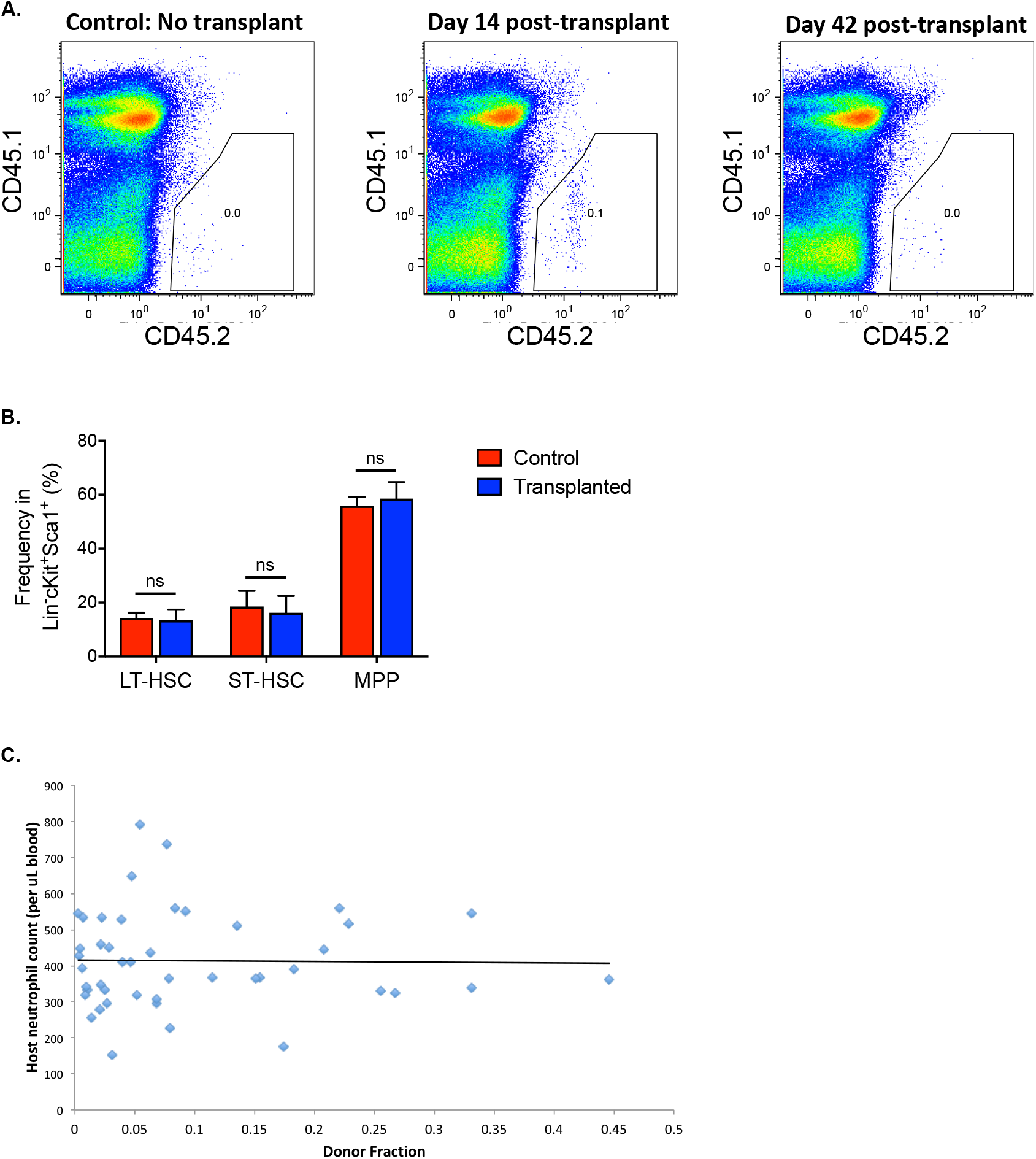
(A) The frequency of donor-derived cells (CD45.2^+^) in whole bone marrow of CD45.1 recipient mice at 2 weeks and 6 weeks after transplantation. (B) Analyses of bone marrow of control mice and those that received HoxB8-conditional progenitor transplantation, as measured at the 6-month time point. Samples were labeled for long-term HSCs (LT-HSC), short-term HSCs (ST-HSC), and multipotent progenitors (MPP) as described in Chorzalska et al. (2018) *Blood*, 132: 2053-2066. (C) Individual measurements of the fraction of donor-derived neutrophils in blood, accrued from serial sampling of CD45.1 mice that received parental or clonal HoxB8-conditional progenitor lines as described in Figure 1D. A linear trendline was projected, indicating a lack of a relationship between donor frequency and absolute neutrophil count in the blood.

**Figure S2.**
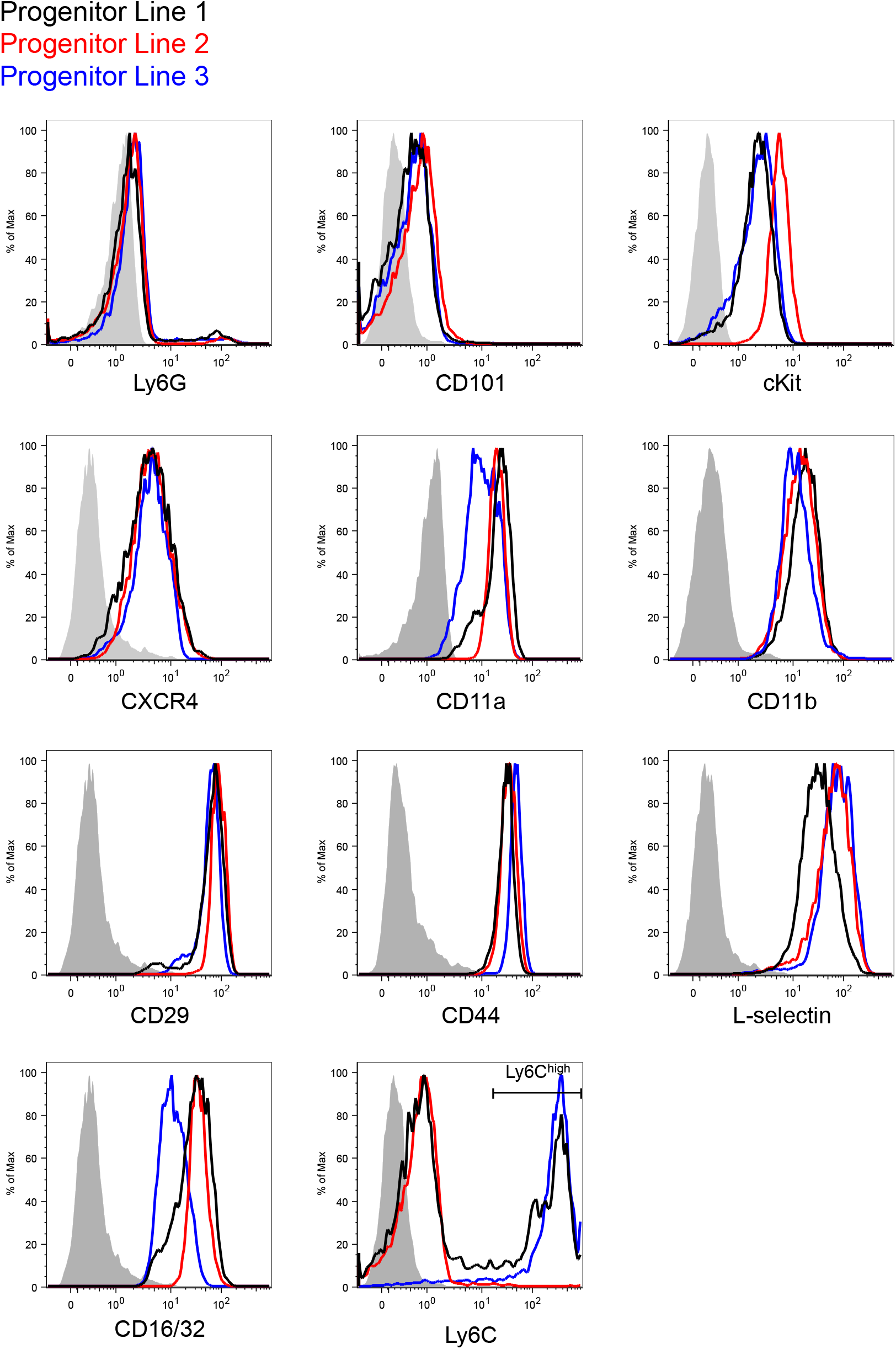
Flow cytometry analyses of the indicated surface receptors and markers on HoxB8-conditional progenitor lines 1, 2, and 3.

**Figure S3.**
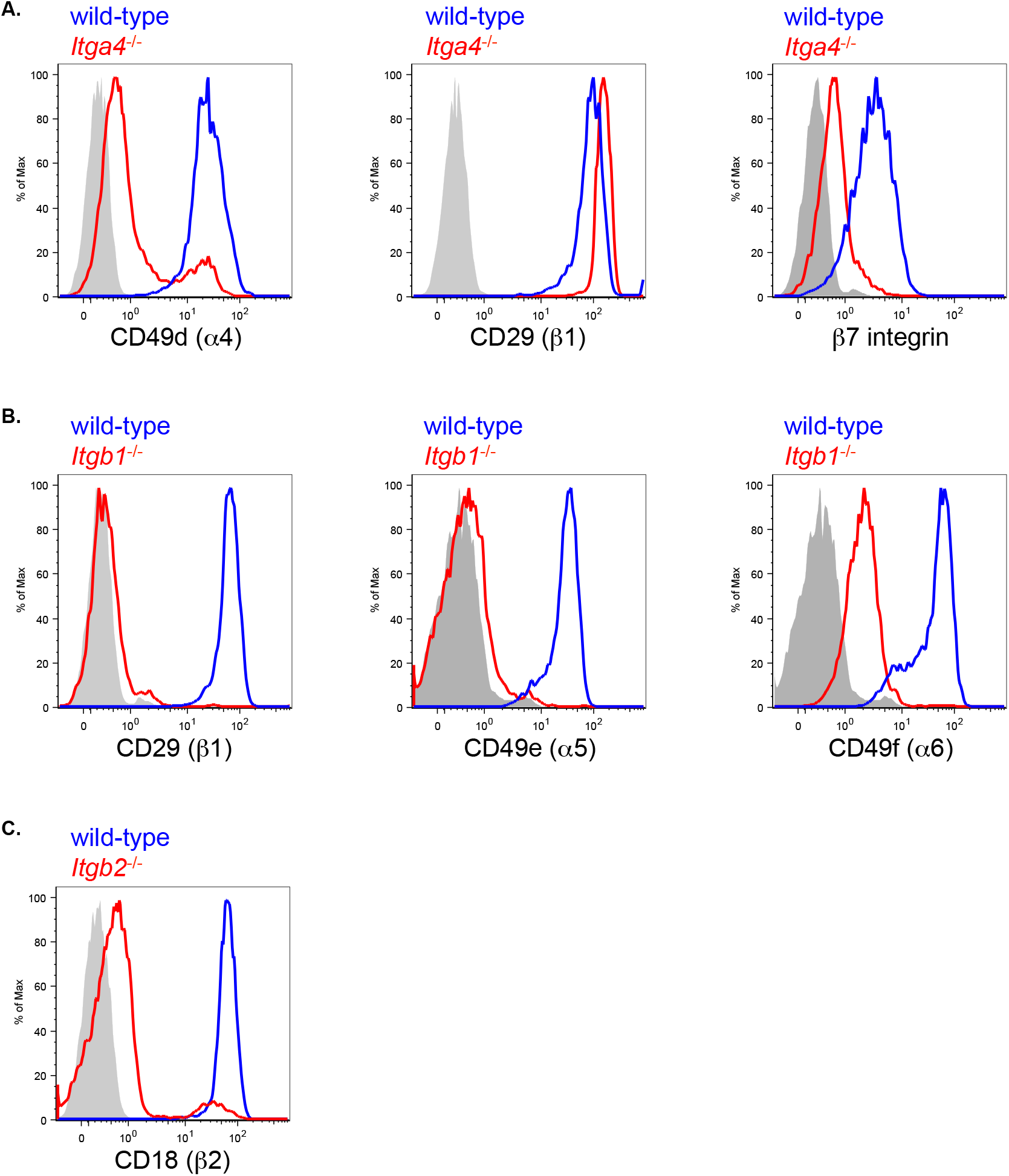
Flow cytometry analyses to determine the expression of the indicated receptors on wild-type and (A) *Itga4*^-/-^, (B) *Itgb1*^-/-^, or (C) *Itgb2*^-/-^ HoxB8-conditional progenitors.

**Figure S4.**
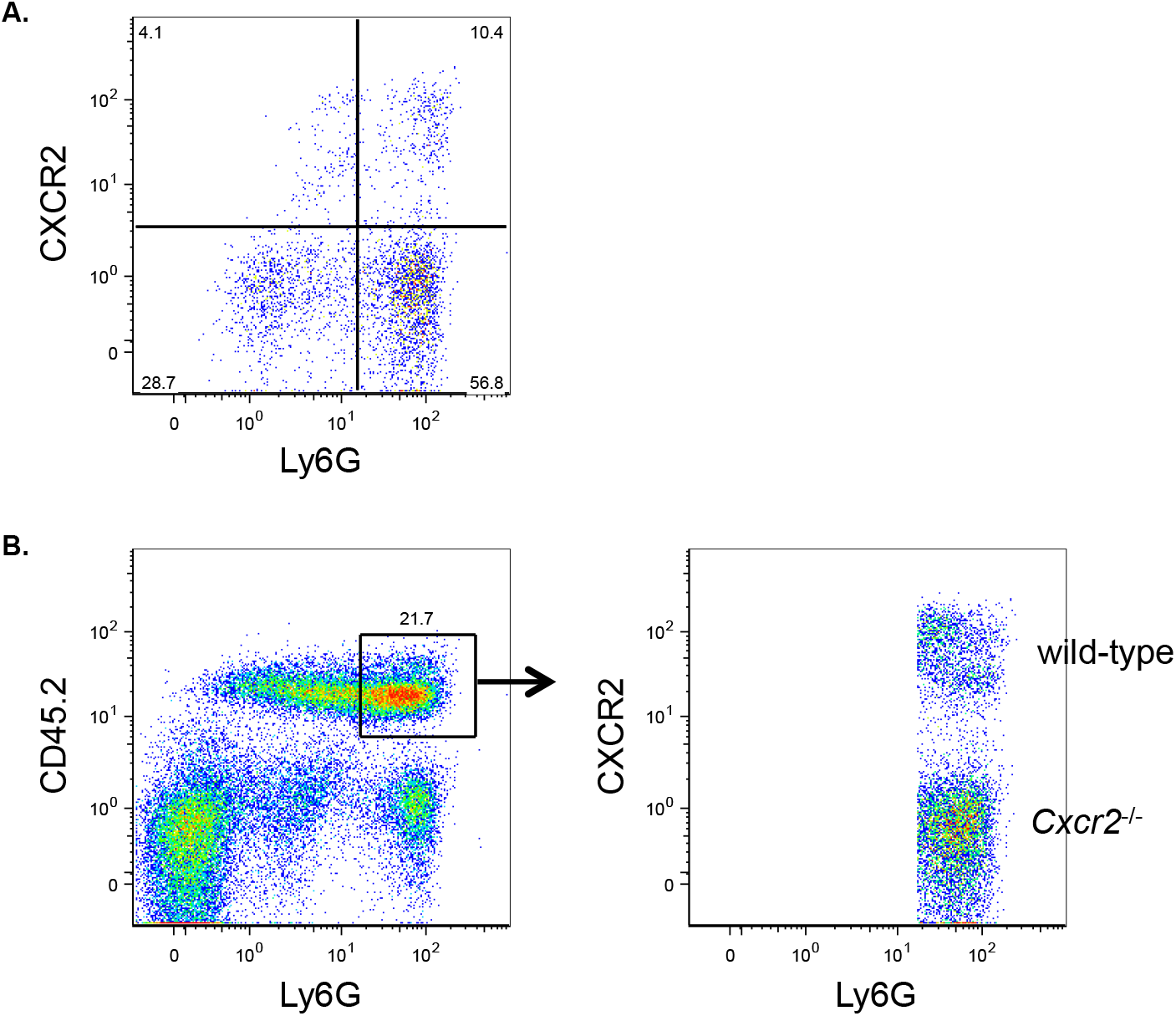
Flow cytometry analysis of donor-derived (CD45.2^+^) cells in the bone marrow of mice showing (A) that *Cxcr2*^-/-^ HoxB8-conditional progenitors are able to differentiate into Ly6G^high^ neutrophils *in vivo*, and (B) the gating strategy for analyses to determine the fraction of wild-type and CXCR2-deficient donor neutrophils within the donor CD45.2^+^Ly6G^high^ population of the bone marrow.

**Figure S5.**
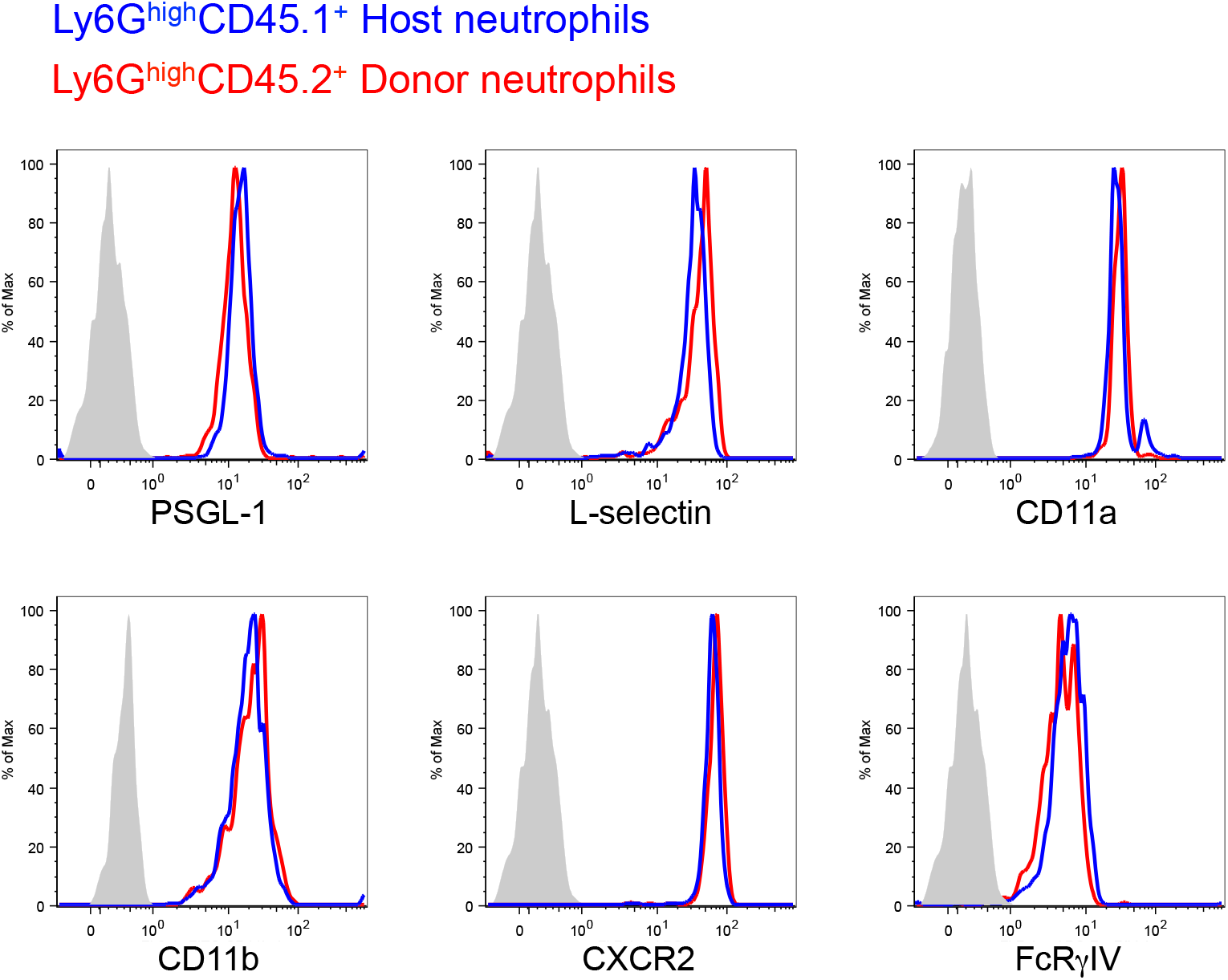
Flow cytometry analyses to determine the expression of the indicated receptors and markers on donor-derived neutrophils (CD45.2^+^Ly6G^high^) in the blood of CD45.1 recipient mice that were transplanted with wild-type HoxB8-conditional progenitors, and their comparison to host-derived neutrophils (CD45.1^+^Ly6G^high^) in the same blood sample.

**Figure S6.**
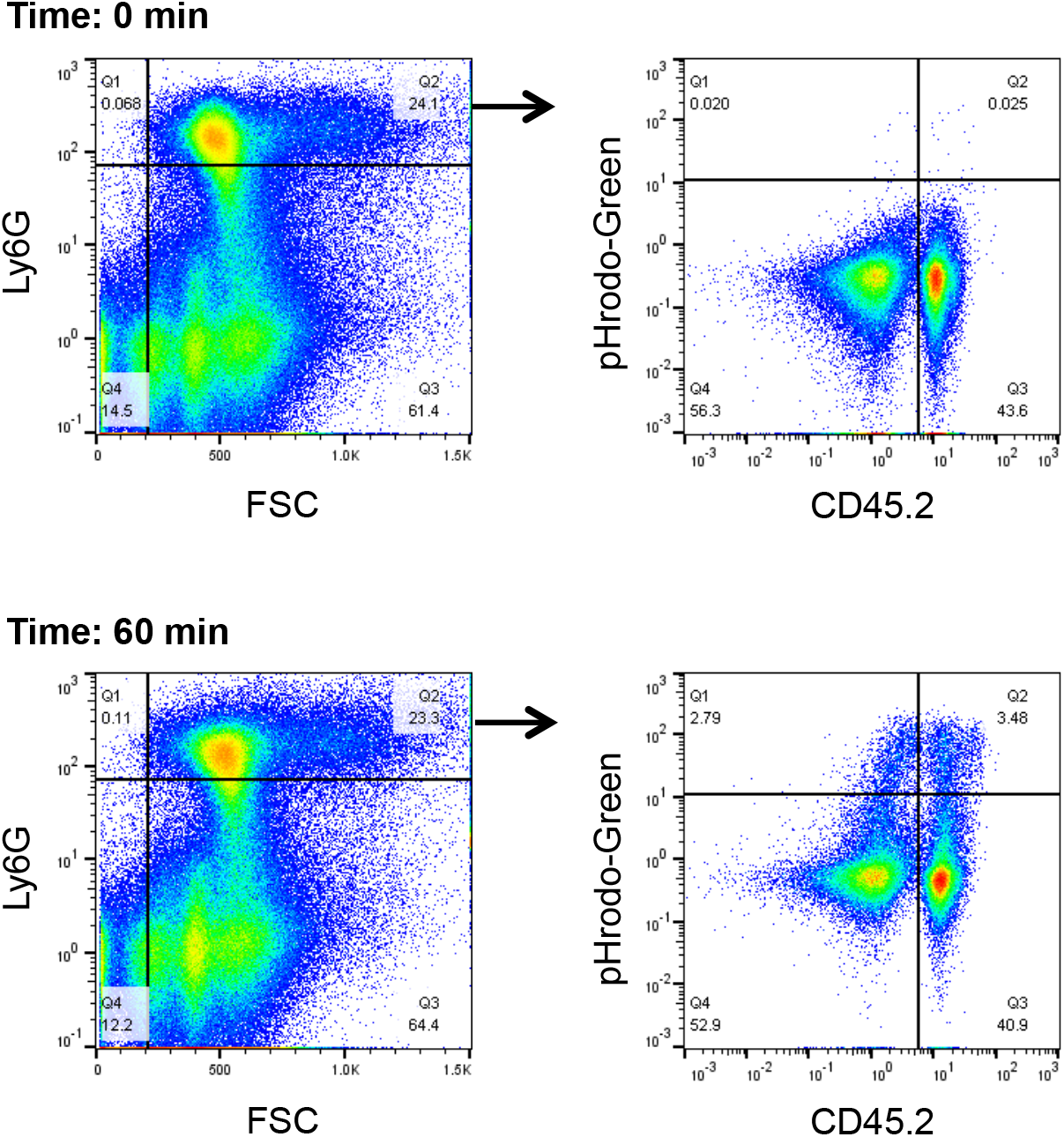
Flow cytometry analyses of phagocytosis assays performed with bone marrow samples. These data indicate the gating strategy to analyze and determine the fraction of donor- and host-derived neutrophils that internalized pHrodo-Green *S. aureus*, at 0 and 60 min time points.

## Notes

### Competing Interest Statement

The authors have declared no competing interest.

